# THE SPLICING FACTOR PTBP1 REPRESSES *TP63 γ* ISOFORM PRODUCTION IN SQUAMOUS CELL CARCINOMA

**DOI:** 10.1101/2021.10.01.462767

**Authors:** William Taylor, David Reboutier, Luc Paillard, Agnès Méreau, Yann Audic

## Abstract

The *TP63* gene encodes the transcription factor p63. It is frequently amplified or overexpressed in squamous cell carcinomas. Owing to alternative splicing, p63 has multiple isoforms called α, β, γ and δ. The regulatory functions of p63 may be isoform-specific. The α isoform inhibits the epithelial to mesenchymal transition (EMT) and controls apoptosis, while the γ isoform promotes EMT. Here, we observed in TCGA data that a high ratio of the *TP63γ* isoform to the other isoforms is a pejorative factor for the survival of patients with head and neck squamous cell carcinoma (HNSCC). We therefore addressed the regulation of the γ isoform. In several tissues (GTEX data), the expression of the RNA-binding protein PTBP1 (polypyrimidine tract binding protein 1) is negatively correlated with the abundance of *TP63γ*. Accordingly, we demonstrated that PTBP1 depletion in HNSCC cell lines leads to an increase in abundance of the γ isoform. By RNA immunoprecipitation and in vitro interaction assays, we showed that PTBP1 directly binds to *TP63* pre-mRNA in close proximity to the *TP63γ*-specific exon. The region around the *TP63γ*-specific exon was sufficient to elicit a PTBP1-dependent regulation of alternative splicing in a splice reporter minigene assay. Finally, we demonstrated that the regulation of *TP63γ* production by PTBP1 is conserved in amphibians, revealing that it encounters a strong evolutionary pressure. Together, these results identify *TP63γ* as a prognostic marker in HNSCC, and identify PTBP1 as a direct negative regulator of its production.

## INTRODUCTION

The *TP63* gene encodes a conserved transcription factor, p63, controlling epithelial development and homeostasis (Mills et al. 1999; Soares et Zhou 2018). In Humans, heterozygous mutations in *TP63* lead to developmental syndromes affecting ectodermal tissue derivatives (Brunner, Hamel, et Bokhoven Hv 2002). Overexpression or amplification of *TP63* is observed in multiple cancer types generally from epithelial origin (Bankhead et al. 2020) and p63 staining therefore serves as a classification biomarker in some malignancies including skin cancer (Smirnov et al. 2019). It discriminates carcinoma from non carcinoma breast cancer types, and lung carcinoma from lung adenocarcinoma in combination with other markers (Travis et al. 2011; Amin et al. 2014). It is also a prognostic marker as *TP63* loss is associated with metastatic progression in head and neck squamous cell carcinoma (Lakshmanachetty et al. 2019).

*TP63* encodes multiple protein isoforms owing to alternative promoters and alternative splicing or polyadenylation. The *TP63* gene structure is evolutionary conserved and beside exons, several intronic regions also present similarities among vertebrates. The complexity of the *TP63* gene structure allows for the production of multiple protein isoforms (**Fig. 1A**). This conserved complexity suggests some evolutionary pressure presumably associated with the function of the different isoforms (Belyi et al. 2010; Lane et al. 2011).

**Figure 1:**
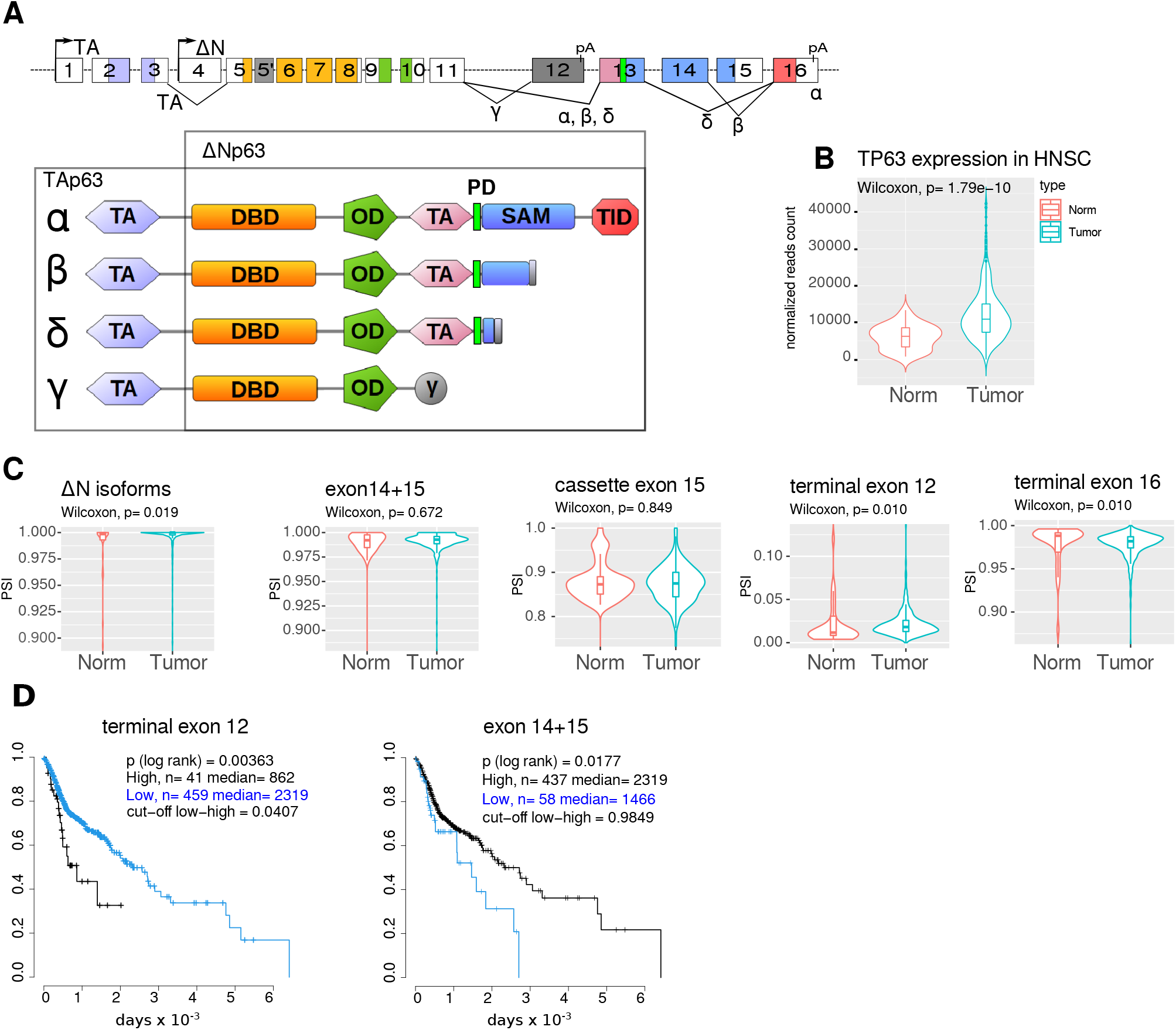
*TP63* splicing from HNSC patients. A) *TP63* exon/intron structure (upper scheme) and corresponding encoded protein isoforms with peptidic domains colour-encoded. TA, ΔN denote the promoters. Exons are numbered according to GTEX. Alternatively spliced junctions and cleavage and polyadenylation events are indicated with solid lines joining exons and pA respectively. Terminal exon 16 is present in isoforms α, β or δ, and terminal exon 12 in isoform γ. The cassette exons 14 and 15 are present in isoform α while the isoform β lacks exon 15 and the isoform δ lacks both. The domains are colored according to the exon/intron structure. TA-Transactivation domain, DBD-DNA Binding Domain, OD-Oligomerisation Domain, SAM-Sterile Alpha Motif, TID-Transactivation Inhibitory Domain, PD-PhosphoDegron. B) Gene-wise expression of *TP63* in normal and tumor samples from HNSC patients. C) Quantification of *TP63* splicing by measuring inclusion of specific exons (percent spliced-in, PSI) in normal and tumor samples. Differential expression between normal and tumor samples was assessed using Wilcoxon rank sum test D) Patients were segregated based on the high or low inclusion of exon 12 (threshold =4.07%) or exon 14+15 (threshold= 98,49 %) in tumor samples. Patient survival was assessed on a Kaplan-Meier graph in the two classes and statistical differences appraised by a log rank test.

The first promoter generates mRNAs encoding TAp63 isoforms comprising the N-terminal transactivation (TA) domain. The mRNAs produced from the second promoter encode only a partial TA domain and the encoded proteins are the ΔNp63 isoforms. The ΔNp63 isoforms are the most abundant ones in epithelial tissues, where they are present mainly in basal cells. They carry most of the p63 functions required for epidermal proliferation and differentiation. The TA isoforms are preferentially expressed during female germline differentiation where they protect the genome from accumulating DNA damage in a way similar to p53 in somatic cells (Suh et al. 2006). A tissue-specific 5’ TA variant coined GTA TP63 has been described in Hominid testis where it is proposed to similarly act as a guardian of the male germ line (Beyer et al. 2011).

Several C-terminal ends are produced from combinations of cassette and alternative terminal exons. These isoforms encode p63 proteins harboring different combinations of the C-terminal domains. The longest isoform p63α contains a phosphodegron (PD) (Galli et al. 2010), a sterile-alpha-motif (SAM) (McGrath et al. 2001), a transcriptional inhibitory domain (Coutandin et al. 2016) and a regulatory sumoylation region (Straub et al. 2010). The shortest mRNA isoform TP63γ encodes only the oligomerisation and DNA binding domains. It is generated by the use of an internal terminal exon and thus has a 3’ untranslated region (3’UTR) different from the other C-terminal isoforms. Since all p63 isoforms contain the oligomerisation domain, they may all interfere with each other’s functions. Most described functions of the α isoform concern its involvement in epithelia, and the p63γ isoform is unable to support proper epithelial development (Wolff et al. 2009). Instead, p63γ promotes the onset of an epithelial to mesenchymal (EMT) transition in keratinocytes (Srivastava et al. 2018), promotes osteoblastic differentiation (Curtis et al. 2015) and favors terminal myogenesis (Cefalù et al. 2015). The functions of p63 are therefore strongly dependent on the regulation of the production of the different 5’ and 3’ isoforms in a tissue-specific manner. Despite the acknowledged importance of the differential role of the p63 C-terminal isoforms, the regulation of the multiple alternative splicing events involved in p63 pre-mRNA maturation remains unaddressed (Pokorná et al. 2021).

Regulation of alternative splicing events requires the assembly of the spliceosome machinery onto splice sites (noted 5’SS and 3’SS with respect to the introns) and the definition of the cleavage and polyadenylation sites (CPA) (Herzel et al. 2017). The tissue-specific choices among different alternative splicing and polyadenylation events are generally controlled by a combination of regulatory sequence elements, silencers or enhancers and their cognate trans-acting factors, generally RNA binding proteins (RBPs). The differential tissue distribution of these RBPs will often account for the tissue-specific regulation of alternative splicing (Fu and Ares Jr 2014). We reasoned that by analyzing relative abundance of RBPs and *TP63* alternative splicing events in RNASeq data from many tissues we could prioritize the RBPs most likely to intervene in TP63 splicing.

Here, we show by combining exploration of TCGA and GTEX Data, RNA/protein biochemistry and reverse genetics experiments that higher inclusion of the γ exon is associated with poorer survival of Head and Neck squamous Cell Carcinoma (HNSCC) patients and we identify the RBP PTBP1 as a direct inhibitor of the inclusion of the γ exon in HNSC cell lines. This regulation of TP63 alternative splicing by PTBP1 appears to be essential as it is evolutionarily conserved, being present in *Xenopus laevis*, a model amphibian that diverged from amniotes about 360 Million years ago. Targeting the PTBP1-mediated regulation of *TP63* expression may therefore be a means to modify TP63 isoform ratio.

## MATERIAL AND METHODS

### Cell lines

HNSC cell lines (TCP-1012) were obtained from ATCC, HaCaT cells were obtained from CLS Cell Lines Service. HeLa, HaCaT, FaDu and Detroit 562 cells were cultured in DMEM medium (Gibco), A253 cells were cultured in Mc Coy’s 5A (Gibco), SCC-9 and SCC-25 cells were cultured in DMEM/F12 (Gibco) supplemented with 400 ng/mL of hydroxycortisone (H0888, Sigma-Aldrich). For siRNA depletion, cells were seeded on 6-well plates at 150000 cells per well. After seeding, a jetPRIME transfection mix (Polyplus) was added, containing 100 pmol of target siRNA or negative controls.

### Xenopus embryos microinjection

Xenopus laevis eggs were obtained from WT females and fertilized using standard procedures (Paris et al. 1988). When indicated 30 ng of MoPtbp1 (Noiret et al. 2016) or 30 ng of control morpholino (GeneTools) was injected into each blastomere of two-cell embryos in a volume of 13.8 nl, using a Nanoject II (Drummond). For rescue experiments, 1 fmol of mRNA-encoding morpholino-resistant PTBP1-V5R was co-injected. Embryos were allowed to develop at 22 °C and collected at stages 26 according to Nieuwkoop and Faber stages (Nieuwkoop et Faber 1994). Total RNA was extracted using RNeasy columns (Qiagen) from a pool of 3 embryos at stage 26. RNA were analysed by RT-PCR for 25 cycle using radiolabeled PCR primer.

### Minigene construction and analysis

The TP63 exon 12 minigene was constucted by gibson assembly of Beta-globin and TP63 gene fragments obtained by PCR amplification from HaCaT genomic DNA using oligonucleotides pairs hGbE1_E2, hTP63_gamma, hGb_E3 and cloned into ApaI, NheI digested pmirglo (Promega). Construction was controled by sequencing. HaCaT cells were transfected with plasmid minigene and selected using G418 (Gibco). Expression and splicing of the minigene was assessed by RT-qPCR using primers presented in the reagent table.

### RNA extraction, reverse transcription and qPCR

Total RNA was extracted from cells using nucleospin RNA kits (Macherey Nagel, #740955.50) following the manufacturer’s instructions. One µg of total RNA was reverse transcribed with superscript II enzyme (Invitrogen) and random primers (Invitrogen #58875). Dilutions of cDNA (1:20) were amplified with PowerSybr Green Master mix (Appliedbiosystems, #4367659) using a Quantstudio (TM) 7 flex real-time PCR 384 well system.

### Western-blot

Cells were washed with PBS 1X and lysed in 2X Laemmli sample buffer (4% SDS, 20% glycerol, 0,125M tris-Hcl pH 6.8, 2.5% β-mercaptoethanol, 0.2% bromophenol blue). Denatured protein samples (10’, 95°C) were loaded onto 8-12% SDS-PAGE gels. Transfer onto nitrocellulose membranes (Amersham Protran 0.45µm NC) was performed using the trans-blot turbo transfer system (Biorad). Membranes were blocked for one hour with a 5% fat-free milk TBST solution (1X TBS, 0.1% Tween 20), incubated with primary antibody solutions (16h, 4°C). After three washes in TBST (5’), the membranes were incubated with anti-mouse or anti-rabbit secondary antibodies coupled to HRP (1h at 20°C), washed three times in TBST (5’). Revelation was performed on an Amersham AI680 imager with either ECL-select (Amersham) or West pico PLUS (Thermo-fisher scientific) chemiluminescent substrate, depending on anticipated signal strength.

### TCGA data analysis

TP63 isoform and splicing in HNSC patient samples were quantified using clinical and expression data from firebrowse.org and splicing quantification from TCGA splice SEQ (Ryan et al. 2016). All expression, splicing and survival analysis were conducted in R. For kaplan-meier estimation, the package survival was used and a log rank test (Bland et Altman 2004) applied. Scripts are available from Gitlab (project ID:29285922).

### GTEX data analysis

To identify RBPs with tissue expression correlated to TP63 tissue specific junction uage, we analysed RNASeq data from GTEx Analysis 8.0. The GTEx data are composed of 17 382 samples from 54 different tissues obtained from 980 donors. For each sample *s*, the total number of reads *Ts* is calculated. The median read depth *Mrd* is calculated form each sample and a read depth normalisation factor (*Ns*) for each sample as:

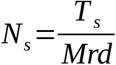

Normalized gene expression *Egs* for gene *g* and sample *s* is computed as:

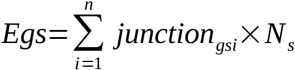

Tissue and samples expressing TP63 are selected and used for correlation analysis. RBPDB (Cook et al. 2011) provided a list of RBPs for which we calculated the RNA expression in the TP63 expressing samples. For each of the 21 TP63 junctions the junction usage is computed as:

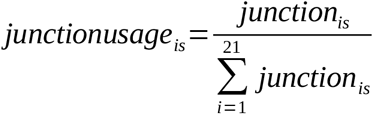

The pearson correlation coefficient and associated p-value is calculated for each RBP with the junction usage of the TP63 junction. Top correlated RBPs are then selected for further analysis. R scripts are available on gitlab (project ID:29285922).

### Fluorescent RNA

Transcription templates were synthesized by PCR (GOTAQ G2, Promega) using a hybrid T7promoter/sequence-specific forward primer and a sequence-specific reverse primer. PCR products were checked on native agarose gel and purified on columns (DC5, Zymoresearch). Fluorescent RNAs were transcribed (37°C, 3h) in (Transc. buffer (1X), 5mM dTT, rNTPs 0.5 mM each, Cy3-UTP 0.1mM (Jena Bioscience), RNAsin (1U/µl, Promega), T7 RNA polymerase (1,25 U/ µl), DNA template (12,5 ng/µl)). RNA were purified on G50 sephadex (GE-healthcare). and controlled by denaturing electrophoresis and fluorescent detection (Typhoon FLA 9500). Cy3 fluorescence and RNA absorbance were quantified on a De Novix DS-11 spectro/fluorimeter.

### Protein production and EMSA

The sequence encoding hsPTBP1 (NM_002819.5) was obtained from sourcebiosciences (IRAUp969B052D). The PTBP1 ORF was PCR amplified and subcloned by gibson cloning into pET21A+ (Novagen) digested by *XhoI* and *NheI*. The 6xHis-tagged hsPTBP1 was expressed in *E*.*Coli* (BL21) after induction by IPTG (1mM), and purified on nickel column using standard procedures. Protein was eluted with 250 mM Imidazole and concentrated on vivaspin (30KDa cutoff). Purifed protein was resuspended in (Sodium Cacodylate pH7.0 20mM, NaCl 100mM, EDTA 0.5 mM, DTT 1mM).

The EMSA experiment were performed by mixing dilutions of hsPTBP1 in RNA Binding buffer (RBB: Sodium Cacodylate pH7.0 10mM, BSA 0.1µg/µl, yeast tRNA 0.1µg/µl, NaCl 100mM, MgCl2 1mM, DTT 1mM, Rnasin (Promega) 0.4U/µl, Heparin (Sigma) 1U/µl) with an equal volume of labeled RNA in RBB. RNA protein complexes were analyzed on native polyacrylamide gels (CLERTE et HALL 2006). Bound and free RNA were quantified on a Typhoon FLA 9500. To estimate the Kd, a non linear model of the form (bound/total RNA) ∼ 1/(1 + Kd/[PTBP1]) was fitted to the data for each RNA using R script (Ryder, Recht, et Williamson 2008).

#### PTBP1 complex immunoprecipitation

Ten millions cells were lysed in (50mM TrisHCl, 0.1M NaCl, 1 % NP40, 0.1 % SDS, 0.5 % Deoxycholate, protease inhibitor cocktail P8340 0.1 %, RNAsin 400U, TurboDNAse 10U, DTT 1mM) and incubated 10 min at 37°C. Prior to immunoprecipitation, lysates were partially digested with RNAseT1 (Ambion) (0.25U per mg of lysate 5 min, 37°C). Lysates were pre-cleared with 50µl of dynabeads Protein-G magnetic for 1 hr at 4°C. The cleared supernatants were incubated with 50 µl of beads pre-coated with 40 µg of anti-PTBP1 antibodies (clone BB7) or mouse IgG Isotype Control (Invitrogen). Immunoprecipitations were conducted overnight at 4°C with constant shaking, then washed successively with IP300 (50 mM HEPES-K pH 7.5, 300 mM KCl, 0.05% NP-40, 0.5 mM DTT), IP500 (50 mM HEPES-K pH 7.5, 500 mM KCl, 0.05% NP-40, 0.5 mM DTT), IP750 (50 mM HEPES-K pH 7.5, 750 mM KCl, 0.05% NP-40, 0.5 mM DTT) and finally once with WB (20mM Tris-HCl pH 7.4; 10mM MgCl2; 0.2%Tween20).

One tenth of the immunoprecipited samples were used for analysis of immunoprecipitated proteins by SDS–PAGE and western blotting. Co-immunoprecipitated RNAs were isolated by proteinase K treatment (30 min, 37°C in 150 µl of PK buffer (50 mM Tris pH 7.4, 1M NaCl, 0.5% NP40, 0.5mM EDTA, 0.1% SDS, 2M Urea, 1% Deoxycholate) containing 50 µg tRNA and 12 U proteinase K (ThermoFisher) followed by phenol–chloroform extraction and ethanol precipitation. RNAs were analyzed by RT–PCR or RT-qPCR.

## RESULTS

### Higher TP63 γ exon inclusion is associated with poor outcome in HNSC patients

While many HNSC tumors overexpress *TP63* (**Fig. 1B** and (Campbell et al. 2018)), the expression and impact of its C-terminal isoforms on patient survival is unclear. We used TCGASpliceSeq data (Ryan et al. 2016) data to evaluate inclusion of exons composing the splice variants of *TP63* in HNSC patients. The *ΔNTP63NTP63* isoforms are characterized by transcription initiation at exon 4 (**Fig.1A and SFig. 1A**) and account for more than 99 % of total *TP63* (**Fig.1C**) in both normal and tumor samples. The *TATP63* isoforms are barely detectable. Among the two terminal exons 12 and 16, the inclusion of exon 16 accounts for about 98 % of the *TP63* RNA (isoforms α, β or δ, see **SFig.1B** for isoform structure). Conversely, usage of exon 12, found exclusively in *TP63γ*, is weak (around 2 % in average). A small but significant increase in the usage of exon 12 is observed in tumors samples compared to normal tissues (p=0.010, Wilcoxon test) with a parallel decrease in exon 16 (**Fig.1C**). The exclusion of exon 15 measures *TP63β*, while the exclusion of the pair of exons 14 and 15 measures *TP63*δ. No difference could be observed between tumors and normal sample for these splicing events. We sought to determine whether differential exon usage in tumors could be associated with difference in survival probability for patients (**Fig.1D**). The measure of inclusion of exon 12 or of both exon 14+15 in tumor samples could discriminate patient survival rates. Higher inclusion (psi >= 4.07%) of the exon 12 was associated with a lower survival of patients (median half-life = 862 days compared to 2319 days, p = 0.00363). Lower inclusion of exon 14+15 was also associated with a more moderate decreased survival (psi < 98.5%, median half-life = 1466 days compared to 2319 days, p = 0.0177). A higher inclusion of *TP63* γ exon appears therefore detrimental for patient survival.

### Candidate RBPs controlling TP63 γ exon inclusion

Because tumors samples may be highly heterogeneous due to tumor type, location or mingled tissues, we sought to identify potential regulators of *TP63* splicing using normal tissues data. We searched for RNA-binding proteins (RBPs) whose expression correlated to *TP63* junctions usages. RNASeq junction quantification were obtained from the GTEX consortium (Lonsdale et al. 2013). They represent quantifications from 54 different tissues or conditions. The correspondence between GTEX and TCGA nomenclatures for *TP63* is shown in SF1A. We calculated the normalized expression for *TP63* in each sample and all tissues. Among the 54 tissues, ten (10/54, 18.5 %) were selected as expressing *TP63* (**Fig.2A**). These tissues where mainly of epithelial origin, as expected for *TP63*, but EBV-transformed lymphocytes and skeletal muscles also showed a notable *TP63* expression.

**Figure 2:**
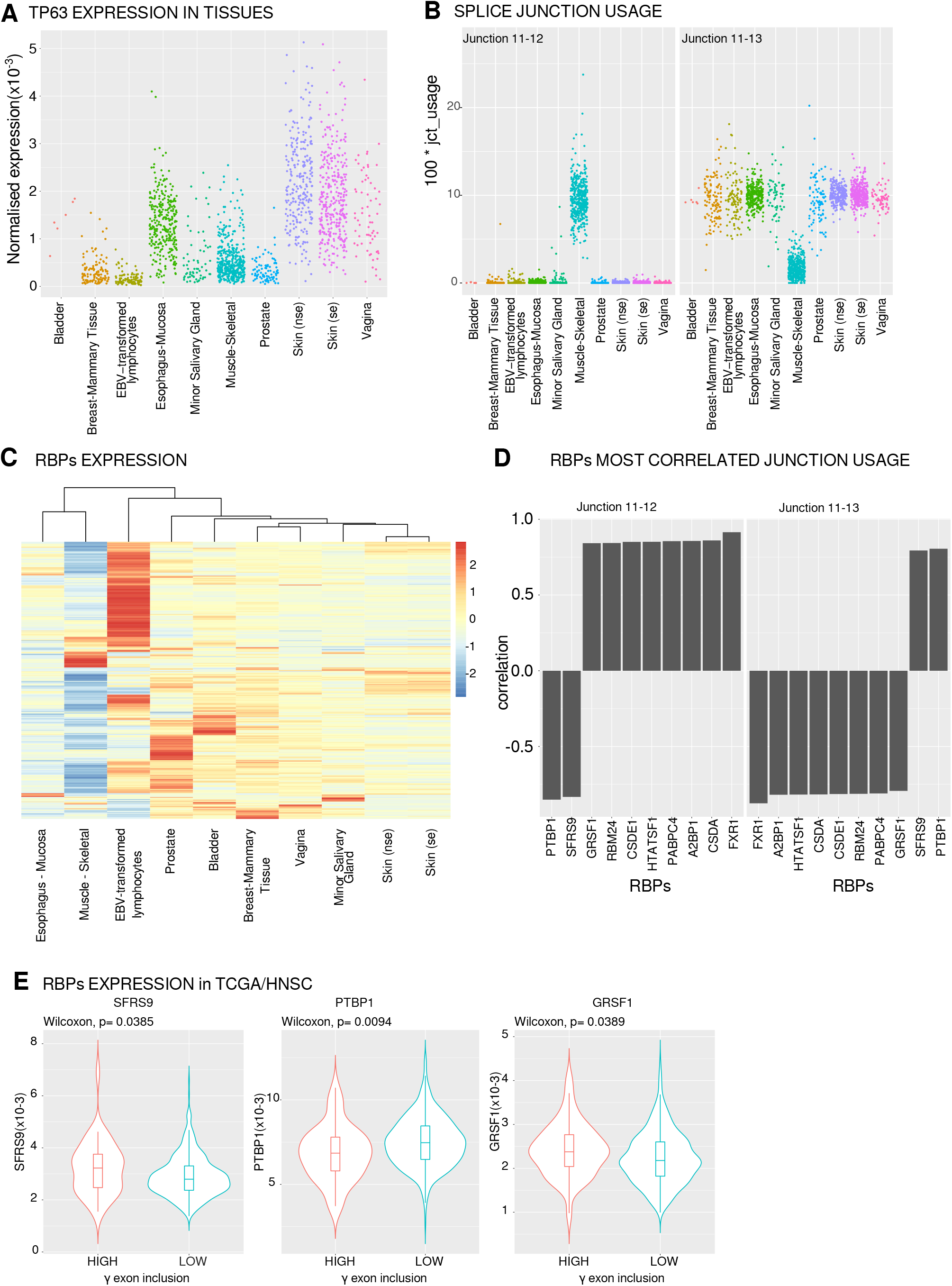
Identification of *PTBP1* as a RBP correlated to *TP63 γ* terminal exon splicing. A) *TP63* gene expression in samples from *TP63* expressing tissues (normalized reads count) (se, nse, (non-) sun exposed). B) Quantification of the usage of junctions 11-12 and 11-13 pertinent to γ terminal exon inclusion. C) RBP expression heatmap and hierarchical clustering of tissues based on RBP expression from GTEX data. D) Top 10 RBPs most correlated to the junctions involved in γ exon inclusion. E) Expression of *SFRS9, PTBP1* and *GRSF1* in two classes of patients with high or low γ exon 12 inclusion (threshold =4.07%) in tumors.

For each tissues and sample we quantified the usage of the 21 individual *TP63* junctions (**SFig.2A)**. The junction usage is strikingly different between tissues (**SFig.2B**). We can distinguish three groups of tissues: Muscle, Epithelia, and EBV-transformed lymphocytes. Eleven junctions are used at similar levels across tissues. Among those, four are very weakly used (J4-6, J5-5’, J5’-6 and J13-16) and seven are constitutive splicing events included at high levels in all tissues. Ten junctions are differentially used across tissues. Junctions (J1-2, J2-3, J3-5, J4-5) are representative of differential promoter usage between epithelial tissue (N isoforms favored) and muscle and EBV-transformed lymphocytes (TA isoforms favored). Muscle specific differences (J11-12 and J11-13, red boxes) are visible. Theses differences are not observed in EBV-transformed lymphocytes. The competition between these two mutually exclusive splicing reactions (J11-12, J11-13) sharing an identical 5’ splice site allows for the production of the γ or [α, β, δ] terminal isoforms. The quantification of these two reactions (**Fig.2B**) clearly shows that they are oppositely regulated, with epithelial tissues favoring the exclusion of the exon γ and the inclusion of the γ terminal exon being promoted in skeletal muscle.

We hypothesized that correlations between RBP expression and junction usage could allow us to pinpoint RBPs acting as splicing enhancers or splicing silencers affecting the tissue specific inclusion of the γ exon. Using the RBPs represented in the RNA Binding Protein Database (Cook et al. 2011), we first determined their expression in *TP63* expressing tissues. We could analyse the expresssion levels of 386 different RBPs in the different tissues (**Fig.2C**). Based on individual samples, we computed the correlations between RBP expression and junction usage relevant to the γ exon (J11-12 and J11-13). Among the 10 most correlated RNA binding proteins (**Fig.2D**), eight were positively correlated and two were negatively correlated to γ exon inclusion (J11-12). As expected the correlation was inverted when considering the alternative junction (J11-13). These 10 RBPs are strongly expressed in muscle (FXR1, GRSF1, RBM24, CSDE1/UNR, HTATSF1, PABPC4, A2BP1/RBFOX1, CSDA/YBX3) or in epithelial tissues (PTBP1, SRSF9).

To determine whether expression of the selected RBPs was coherent with higher γ exon inclusion in HNSC patients with poor survival, we compared the differential expression of the RBPs in tumors selected for having high or low levels of γ exon that discriminate among patient survival (**Fig.1C**). Of the 10 RBPs tested, only three (SFRS9, PTBP1, GRSF1) are differentially (Wilcoxon, p<0.05) expressed at the RNA level between tumors from patients with high or low γ exon inclusion (**Fig.2E**). *SFRS9* is more highly expressed in samples with high γ exon inclusion (Wilcoxon, p= 0.0385), which is not in accordance with the GTEX data where *SRSF9* is negatively correlated to γ exon inclusion (**Fig.2D**). *PTBP1* and *GRSF1* are low and high, respectively, in patients with high gamma (Wilcoxon, p= 0.0094 and p= 0.0389, **Fig.2E**). The same RBPs are negatively and positively correlated, respectively, with γ in GTEX data (**Fig.2D**). Therefore PTBP1 and GRSF1 can coherently be involved in γ exon regulation. However, because GRSF1 is mainly a mitochondrial RBP and is weakly expressed in epithelial cells (Jourdain et al. 2013) while PTBP1 is a well defined alternative splicing regulator expressed in epithelial tissues, we chose to test whether experimental depletion of PTBP1 could enhance *TP63* γ exon inclusion in a panel of HNSCC cell lines (FaDu, SCC9, SCC25, Detroit562 and A253) from different tissues of origin.

### PTBP1 represses endogenous TP63 γ exon inclusion in squamous carcinoma cell lines

Using two different antibodies we analyzed p63 and p63α-specific expression by western-blot (**Fig.3A**) in a panel of HNSC cell lines. Both antibodies detected a major isoform around 70 kDa, coherent with the major ΔNP63α isoform (MW 66 kDa). In addition to the 70 KDa protein the pan-p63 antibody (4A4) detected lower molecular weight bands undetected using the α-specific antibody (D2K8X). This suggested that isoforms other than ΔNP63α are expressed at lower levels in the tested cells. The identity of these isoforms can not be inferred from these experiments. We used RT-qPCR to evaluate the abundance at the RNA level of total *TP63* and of its three predominant isoforms (α, β, γ). We estimated that global *TP63* expression levels were higher in SCC25, then A253 and SCC9, then FaDu, Detroit 562 and non transformed keratinocytes HaCaT. As expected, it is barely detectable in HeLa cells used as a negative control (**Fig. 3B**). We used isoform-specific RT-qPCR to quantify C-terminal isoforms (**Fig.3C**). In all cell types tested, the α isoform is the major one ranging from 87 to 95 % of *TP63* mRNA. Detroit 562 cells have the lowest α proportion. Accordingly, β and γ isoforms are in higher proportions in Detroit 562 cells with 8 % and 2.7 % of each respectively. The expression of the β and γ isoforms are variable among the cell lines with the Detroit 562 and the SCC25 cells having the highest relative proportion of TP63γ (2.75 and 1.5% respectively) while other cell lines had all less than 1 % of TP63γ mRNA (**Fig.3C**).

**Figure 3:**
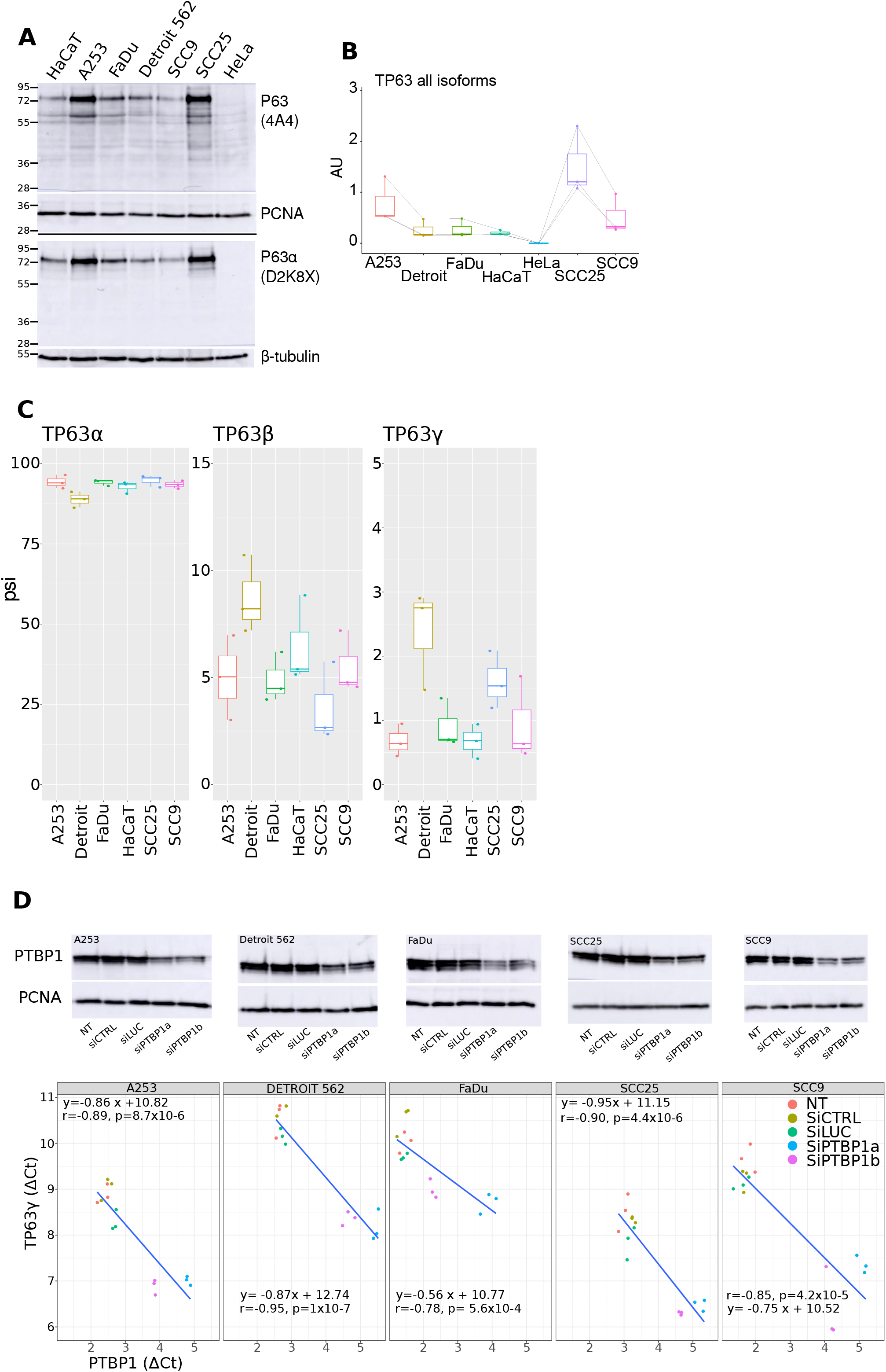
PTBP1 controls TP63 splicing in HNSCC cell lines. A) p63 expression was evaluated in HaCaT, HeLa and HNSC cells by western blot and immunodetection using a pan-p63 antibody (4A4) or a p63α-specific antibody (D2K8X) (n=3). Even loading was controlled using PCNA or β-tubulin antibodies as indicated. B) *TP63* RNA expression was measured by RT-QPCR with one pair of primer detecting all isoforms and quantification was normalised to a calibration curve obtained from TP63 plasmid DNA dilutions (n=3). C) Quantification of *TP63*α, β or γ percent spliced-in using isoform-specific primer pairs (n=3). D) upper panel, quantification of *TP63*γ isoform and *PTBP1* RNA in control and PTBP1 depleted HNSC cell lines (n=3). Lower panel, evaluation of PTBP1 depletion by western blot (n=3), PCNA serves as a loading control.

We efficiently depleted PTBP1 in the five HNSC cell lines using two different siRNAs (**Fig.3D**, upper panels). In all five cell lines, depletion of PTBP1 led to an increase in *TP63γ* abundance as shown by the lower *TP63γ* ACT in PTBP1 depleted cells (**Fig.3D**, lower panels). The *TP63γ* levels were therefore inversely related to PTBP1 levels. This demonstrates that PTBP1 represses the accumulation of the *TP63γ* isoform in HNSC cell lines.

### PTBP1 binds to TP63 pre-mRNAs sequences in vivo on the *γ exon* 3’Splice site and CPAs

Direct regulation of *TP63* splicing would require binding of PTBP1 to regulatory elements located in the pre-mRNA of interest. To determine whether PTBP1 was bound to *TP63* pre-mRNAs *in vivo* we immuno-precipitated specifically PTBP1/RNA complexes from HaCaT extracts and analyzed the pre-mRNA content of the complexes by RT-PCR and RT-qPCR (**Fig.4A, 4B)**. Presence of *TP63* pre-mRNA was specifically detected in input and PTBP1 immunoprecipitates but not using control Ig or beads only. To determine whether PTBP1 was preferentially associated to *TP63* pre-mRNA or mRNA, we used primers specific to *TP63* mRNAs (ex8-ex9, ex11-12) or pre-mRNA (int10-ex11, int11-ex12, int12-ex13). As shown in **Fig.4B**, the most significant and robust enrichment was observed with the primer pair specific to the pre-mRNA region closest to the γ exon (int11-ex12). Control RT-qPCR performed in absence of reverse transcriptase did not show PCR amplification (data not shown). Therefore, on *TP63* pre-mRNA, PTBP1 is preferentially bound to the region that is up-regulated following PTBP1 depletion.

**Figure 4:**
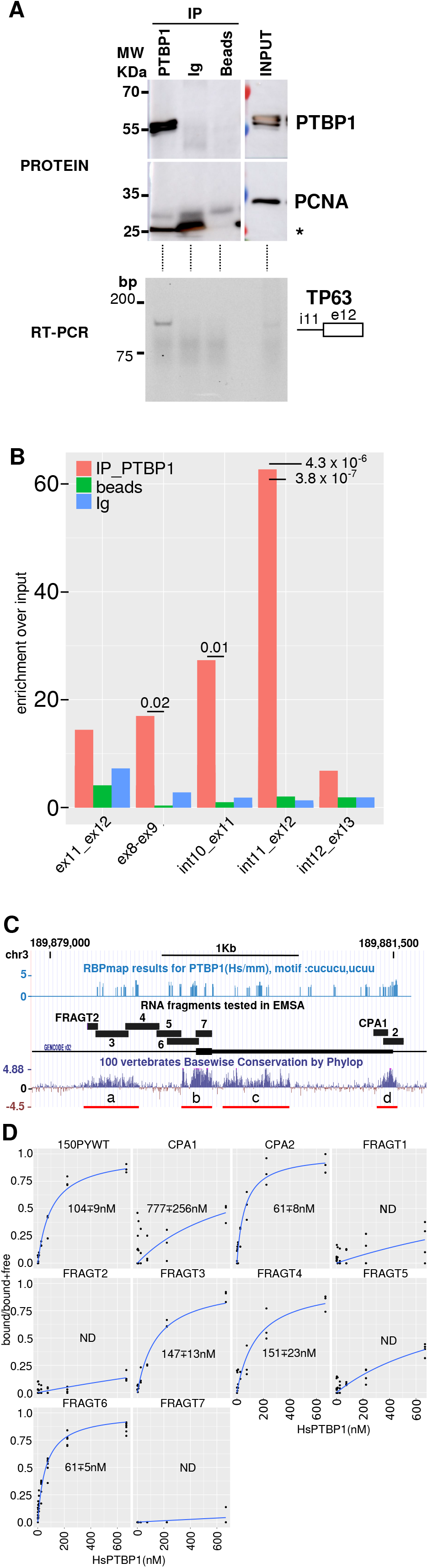
PTBP1 binds to TP63 premRNAs sequences. A) Immuno-purification of PTBP1/RNA complexes or control experiments (Ig, beads only) from HaCaT cell extracts. Immunopurification was controlled by PTBP1 and PCNA western-blot as indicated in input (10%) or IP samples (upper panels). Protein MW are indicated on the side of the membranes. The asterisk denotes the light-chain of the IgG. Lower panel, RT-PCR detection of *TP63* pre-mRNA with primers located in intron 11 and exon 12 in the indicated samples. B) RT-qPCR detection of different mRNA or pre-mRNA portions of *TP63* from IP or control samples relative to input signal. Statistical assessment between PTBP1 and control IP was measured by a student-test (n=3). C) *TP63* exon 12 genomic locus and position of the RNAs tested in RNA/protein shift assays. Upper lane, RBPmap predicted PTBP1 binding sites. Middle lane, positions and names of the RNA fragment tested along the *TP63* exon 12 genomic sequence. Lower lane, nucleotidic conservation among vertebrates with regions of high conservation labeled a,b,c,d as discussed in the text. D) PTBP1/RNAs binding curves (bound/bound + free) obtained for the indicated RNAs (n>=4). The calculated Kd and their s.d. is shown for each plot.

The RBPmap database details known binding data for hundred of RBPs (Paz et al. 2014). Binding motifs for PTBP1 represent predicted binding sites (PBS) and were plotted along the TP63 γ exonic region (**Fig.4C**). Examination of the genomic sequences surrounding *TP63* γ exon identifies 4 blocks of conservation among vertebrates : an upstream intronic region (a), a region overlapping the 3’ splice site (b), a broader area in the γ 3’UTR (c) and a smaller region overlapping the cleavage and polyadenylation (CPA) site (d).

We performed electrophoretic mobility shift assay (EMSA) using recombinant PTBP1 and fluorescently labelled RNA molecules to determine the affinity of PTBP1 for RNA sequences representing the conserved sequence elements (**Fig.4D** and **SFig.3)**. We quantified the fluorescent RNAs in bound and unbound PTBP1 complexes and calculated a Kd based on PTBP1 concentration (Ryder, Recht, et Williamson 2008). As presented in **Fig.4D**, RNA fragment 1 (FRAGT 1) composed solely of β-globin RNA sequences, FRAGT2 that is composed in part of β-globin RNA sequences and in part of the *TP63* intronic region, FRAGT 5, FRAGT7 located totally in the γ exon, and CPA1 located close to the cleavage and polyadenylation site, remain unbound to PTBP1. FRAGT3, FRAGT4 and FRAGT6 and CPA2 associate strongly with human PTBP1 protein with binding affinities better than (CPA2, FRAGT6, ∼ 60 nM) or comparable to (FRAGT3, FRAGT4, ∼ 150 nM) the positive PTBP1binding control RNA (150PYWT, ∼ 104 nM) (Hamon et al. 2004).

Altogether, this demonstrated the direct association of PTBP1 on regulatory regions surrounding the γ exon 3’Splice site and CPAs on the TP63 pre-mRNA *in vivo* in an epithelial cell lines.

### PTBP1 represses TP63 γ exon inclusion in a minigene in epithelial cells

If PTBP1 can control *TP63γ* exon inclusion by directly binding to these cis-elements, then the splicing of a minigene composed of the *TP63* genomic sequence should be under the control of PTBP1. To test this, we stably transfected HaCaT cells with minigenes containing *TP63* genomic sequences within an efficient *β-globin* splicing context. The structure of the minigene is shown in **Fig.5A**. We depleted PTBP1 in HaCaT keratinocytes using either of two different siRNAs (siP1A and siP1B, **Fig.5B**). As previously shown in HNSC cell lines (**Fig.3D**), PTBP1 depletion strongly increases the abundance of TP63 γ in HaCaT cells (**Fig.5C**). This demonstrates that, like in transformed cells, the accumulation of the endogenous *TP63γ* isoform is at least partly repressed by PTBP1 in non-transformed HaCaT keratinocytes.

**Figure 5:**
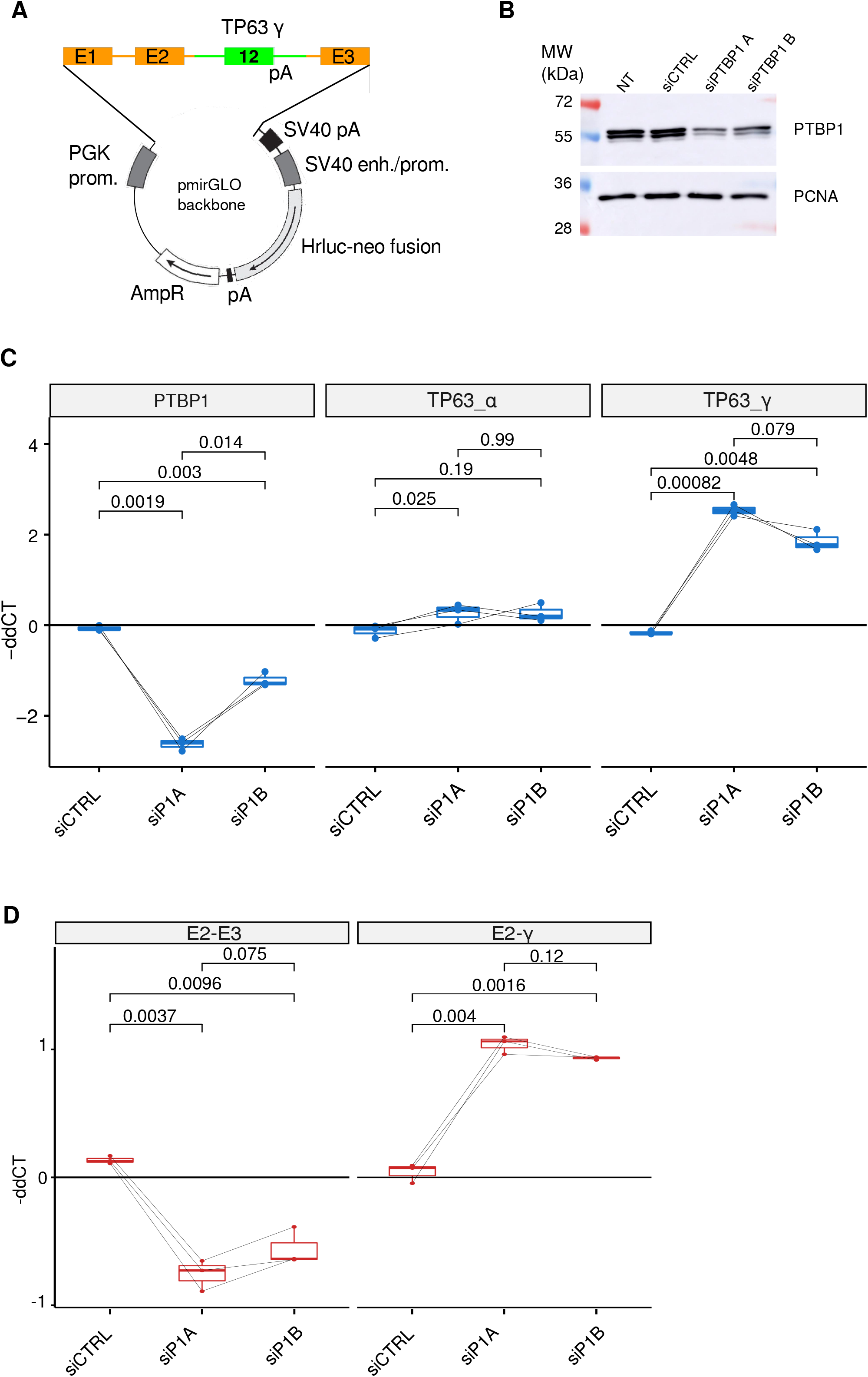
PTBP1 represses endogenous *TP63* γ exon inclusion in HaCaT cells and on a *TP63* γ exon minigene. A) Structure of the *TP63* minigene stably transfected in HaCaT cells. B-globin gene sequence are shown in orange, *TP63* gene sequences are shown in green. B) Detection of PTBP1 in HaCaT cells after PTBP1 depletion mediated by two different siRNA (n=3). C) RT-qPCR quantification of *PTBP1* levels and endogenous *TP63*α and γ isoforms after siRNA-mediated PTBP1 depletion (n=3, t-test). D) RT-qPCR quantification of the RNA isoforms produced from the *TP63* minigene (n=3, t test).

The *TP63* minigene splicing is also dependent on PTBP1 abundance (**Fig.5D)**. Upon PTBP1 depletion, the splicing between E2 and the γ terminal exon was increased 2 times (paired t.test, pvalue = 0.0016, n=3) while the alternative possibility, splicing of E2 to E3, was decreased about 2 times (paired t.test, pvalue = 0.004, n=3). This demonstrates that the genomic region surrounding the γ terminal exon is sufficient to allow for its PTBP1-dependent regulation.

### PTBP1 represses TP63 γ inclusion since at least 350 Millions years

Clusters of PTBP1 PBS are located in-between evolutionarily conserved regions a and b and in regions b, c and d (**Fig.4C**). Because of this interesting conservation of non coding regions, we addressed the regulation of the PTBP1-dependent regulation of the γ exon in an evolutionary distant organism. It is estimated that amphibians and mammals diverged from their common tetrapod ancestor roughly 360 millions years ago (Hellsten et al. 2010). Within amphibians, *Xenopus laevis* is a well-established model of vertebrate development. The exon/intron structure of the *TP63* locus is well conserved between Xenopus and human with conserved coding exons (in blue on **Fig.6A**) serving as milestones along the sequence. In *Xenopus*, two annotated additional exons (11’ and *, in green) are present (**Fig.6A**). Exon * is very weakly used and exon 11’ encode an alternative peptide in frame with the rest of the coding sequence. A terminal exon corresponding to the human gamma exon is present in both species.

**Figure 6:**
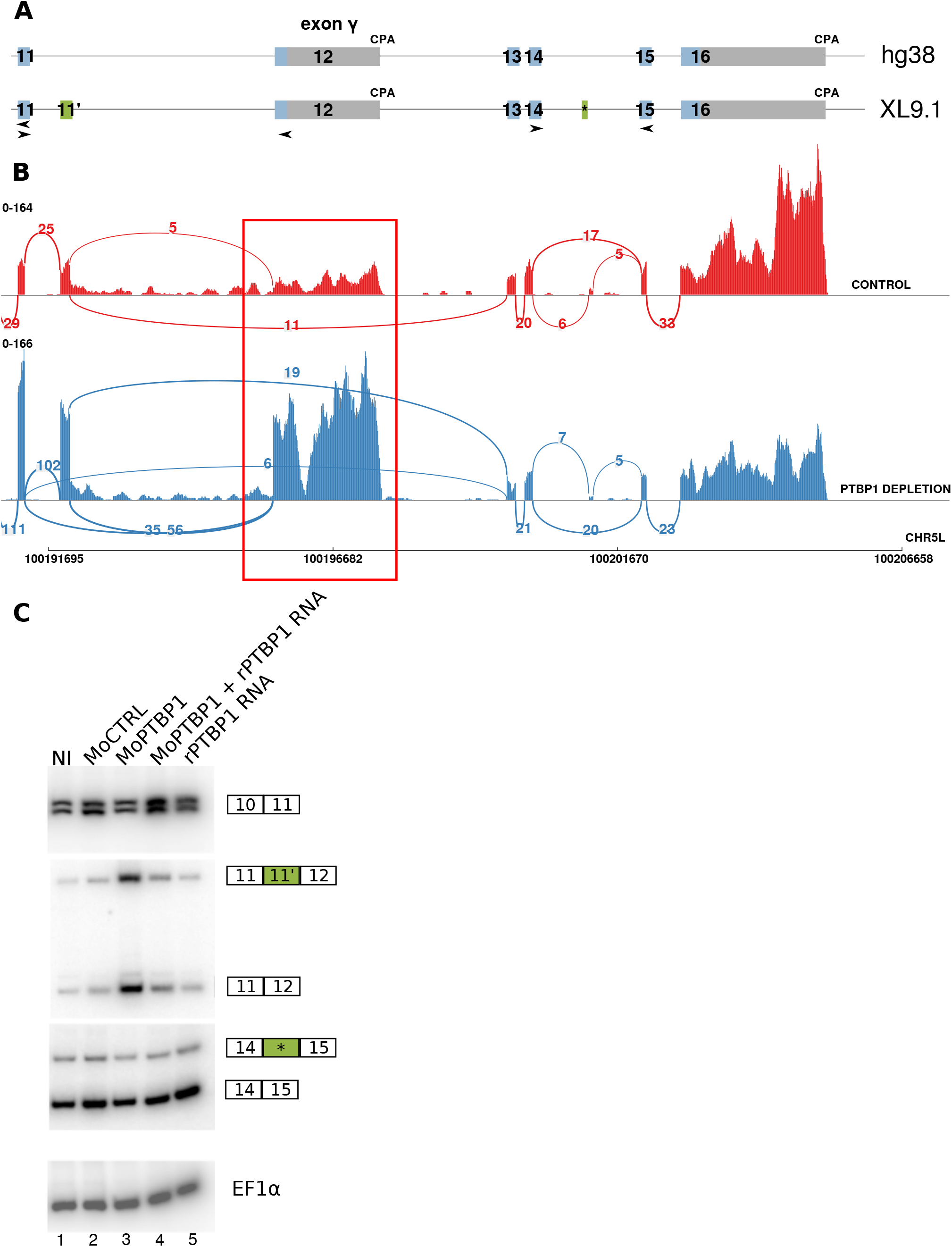
PTBP1 repression of endogenous *TP63* γ exon inclusion is conserved from amphibian to human. A) comparison of gene structure and exon conservation between human (hg38) and *Xenopus laevis* (XL9.1). Conserved coding region are in blue. Frog specific coding exons are in green. Arrowheads indicates primer pairs used in the RT-PCR shown in C. B) Sashimi plot representing the accumulation of mapped reads on the locus of the *TP63*.*L* gene in one control sample (red) and one PTBP1 depleted samples (blue) (n=3). The region undergoing a major shift in mRNA splicing is framed in red. C) RT-PCR analysis of TP63 splicing using isoform specific primer pairs. The RT-PCR products are schematized on the side of the gel. RT-PCR on *EF1α* serves as a control. NI : uninjected embryos, MoCTRL: control morpholino oligonucleotide, MoPTBP1: PTBP1 depleting morpholino, MoPTBP1 + RNA PTBP1: rescue experiment with re-expression of PTBP1 from a Morpholino-immune injected *rPTBP1* mRNA, RNA rPTBP1: Morpholino-immune rescue *PTBP1* mRNA.

Previously, we used antisense morpholino oligonucleotides to deplete Ptbp1 from developing embryos, and we analyzed RNA expression by RNAseq (Noiret et al. 2016). Here, we re-investigated these data to focus on Xenopus *tp63*. Notably, the depletion of Ptbp1 led to an increase in the γ exon (12) inclusion in embryos as judged by the Sashimi plot on **Fig.6B**. On this example, reads spanning junction 11-12 or junction 11’-12 increased from 0 to 35, and from 5 to 56 respectively. This was accompanied by an accumulation of sequencing reads on the γ exon. To independently confirm these data, we analyzed Xenopus embryos by RT-PCR **(Fig.6C)**. The analyzed embryos were previously injected with a control antisense morpholino oligonucleotide (lane 2), a morpholino against PTBP1 (lane 3), the same morpholino co-injected with an mRNA encoding PTBP1 made immune to the morpholino by a few nucleotidic substitutions (lane 4), or the rescue RNA alone (lane 5). As shown on **Fig.6C**, depletion of PTBP1 leads to the specific accumulation of RT-PCR products 11-12 and 11-11’-12, revealing an increased inclusion of the γ exon. That this increase is due to a specific change in RNA splicing in this region is shown by the unchanged signal when another region of *tp63* mRNA is amplified (primers in exons 14 and 15). Upon restoration of PTBP1 expression from an injected rPTBP1 mRNA which is resistant to morpholino inhibition, the γ exon inclusion was restored to control levels.

These results demonstrate that the repression of γ exon usage by the RNA binding protein PTBP1 is a mechanism that emerged at least in tetrapods and has been conserved across 360 Millions years of evolution. Together, these data reveal that PTBP1 controls TP63γ exon inclusion by binding to discrete and conserved regulatory elements located in this pre-mRNA region. The γ exon inclusion is hence restricted by the presence of PTBP1, decreasing the abundance of the TP63 γ isoform in normal and cancerous epithelial cell lines where PTBP1 is abundantly expressed.

## DISCUSSION

Alternative splicing misregulation in cancer cells is emerging as a major theme in the alteration of gene expression observed during the cancerous process. This offer new routes of tumor classification or treatment (Bonnal, López-Oreja, et Valcárcel 2020). Using a heuristic approach we identified *PTBP1* as a major repressor of *TP63γ* production in normal and malignant epithelial cells. This is the first identification of a trans-acting factors controlling the splicing pattern of *TP63* in a tissue-specific manner.

Predicted PTBP1 binding sites were located in evolutionarily conserved regions framing exon 12 in the *TP63* pre-mRNA and we demonstrated that PTBP1 binds with high affinity to discrete elements of the *TP63* pre-mRNAs *in vitro* and in cells. These elements coincide with the 3’ SS and the CPA of the γ exon.

### Model of γ terminal exon regulation by PTBP1

PTBP1 is generally observed as an inhibitor of the splicing or polyadenylation reaction. However some cases of exon activation by PTBP1 have been reported (Lou et al. 1999). This inhibition is generally the consequence of the inhibition of the recognition of the 3’ splice site by U2AF (Saulière et al. 2006) or of the CPA site by CSTF (Castelo-Branco et al. 2004). On X*enopus* tropomyosin mRNA, PTBP1 inhibits the function of an intronic silencer element located upstream of a terminal exon (Le Sommer et al. 2005). The multiple RNA-recognition motifs of PTBP1 and its possible multimerisation on its targets offer multiple modes of interaction (Oberstrass et al. 2005). PTBP1 binding to a pre-mRNA at several sites can either hinder the regulated exon or exclude it by creating a loop through interaction of PTBP1 in the upstream 5’ and downstream 3’ region surrounding the regulated exon (Spellman et Smith 2006). Gruber and colleagues (Gruber et al. 2018) identified repeats of CU dinucleotides, reminiscent of PTBP1 binding sites, as preferentially associated with differentially used CPA in tumors versus normal samples of gliobastomas. *PTBP1* levels were strongly anti-correlated with the length of these RNAs suggesting that PTBP1 acts as an inhibitor of CPA on tandem UTRs harboring sequential polyadenylation sites. Binding of PTBP1 on the CU-rich regions around the CPA would hinder its recognition by the CPA-bound machinery favoring the alternative unmasked CPA. The context of the *TP63* γ exon is different as it is an internal terminal exon in competition with the splicing of a classic exon defined by 5’ and 3’SS. As interactions between splicing factors involved in the recognition of the 3’SS and the CPA bound factors are central in the efficient processing of terminal exon (Millevoi et al. 2006), we can propose that by its dual binding to the 3’SS and the CPA, PTBP1 is likely interfering with the ability of U2AF and CPA factors to interact with 3’SS and CPA respectively and therefore impede their ability to define the γ exon.

In HNSCC cells depleted in PTBP1, the inclusion of the γ exon is still modest (see **Fig.3D**). It is therefore possible that additional inhibitory factors are at play in theses cell lines. We can exclude that this limited effect is because of a redundant role of the orthologues of PTBP1 as co-depletion experiments of PTBP1 and PTBP2 barely increase the γ exon inclusion (data not shown). One alternative is that the limited γ exon inclusion in HNSC cells is due to intrinsic properties of this γ exon, with sub-efficient 3’SS and CPA that would only drive limited inclusion. In this case efficient terminal exon γ inclusion observed in other tissues would require specific factors to enhance exon γ inclusion. Such factors should be looked for in tissues where the exon γ is predominantly used and we can assume that germ cell or muscle cells may prove useful to identify such factors.

### Which role for the γ terminal exon ?

The importance of TP63 in development is clearly demonstrated from works in numerous models. In Xenopus, *tp63* is required for the proper muco-ciliary epithelial development (Haas et al. 2019). The evolutionarily conserved regulation of the γ exon by PTBP1 suggests that the p63γ harbors specific and required functions in certain cellular type or tissues. These functions may come from its impact on the p63 coding sequence, from a difference in the sequences of the 3’UTRs between γ and the α or β isoforms, or from more complex phenomenon such as regulation of circular RNA production as already described for the *TP63* gene (Cheng et al. 2019). At the RNA level, because of its different 3’UTR, and the γ isoforms have a different susceptibility to miRNAs controlling TP63 expression than the α, β isoforms (Lin et al. 2015). It is therefore possible that the production of the γ isoform provides the cell with a way to escape some miRNA-mediated tissue-specific regulation of *TP63*. However this change in 3’UTR would also result in a change in the encoded p63 isoforms.

It is unlikely that the γ-specific peptide encoded in the terminal exon has any conserved functions by itself because of its important divergence between species. Indeed, only 15/37 (40.5%) aminoacids are conserved between Xenopus and Human γ peptides while the overall identity of ΔNp63α proteins is 85 %. Both in Human and in *Xenopus*, the γ isoform is strikingly different from the α isoform. In Human, it is deprived of 231 amino acids forming the SAM domain, a transcription inhibitory domain, a transcription activation domain and a phosphodegron. These domains are of paramount importance for the correct function of TP63 as demonstrated by the Ankyloblepharon Ectodermal defects Cleft lip/palate (AEC) and Rapp–Hodgkin (RHS) syndromes associated with mutations altering these domains (Rinne, Brunner, et van Bokhoven 2007). Yet p63γ retains the DNA binding and oligomerisation domains allowing the protein to form hetero-complexes that have altered transcriptional functionality (Petitjean et al. 2008).

In general, the ability of an alternative isoform to affect protein complex functions increases with the number of monomers composing the complex (Bergendahl et al. 2019). p63 forms homo- or hetero-tetramers composed of various p63 or p73 proteins. It is therefore possible that even low amounts of the γ isoform affect the functions of p63 as a transcription factor. It is unclear yet if the low amount of TP63γ at the RNA levels are strictly correlated to similarly low protein level. Indeed, expression experiments showed that ΔNp63γ was the most stable p63 isoform in Hep3B cells (Petitjean et al. 2008). This may be due to ΔNp63γ being deprived of the Fbw7 E3 ligase phosphodegron and of some of the lysine residues (K494,/505) involved in ubiquitinylation and destabilisation of p63α (Galli et al. 2010; Prieto Garcia et al. 2020). Quantification of the γ isoform at the protein level is complex as no γ -specific antibodies are currently available. We may expect that state of the art mass-spectroscopy will allow for isoform specific quantification of protein in the near future (Lau et al. 2019).

The overexpression of ΔNp63γ in the context of global p63 depletion leads to increased EMT in both a breast cancer cell line (Lindsay et al. 2011) and in a 3D organotypic model of invasion ((Srivastava et al. 2018). This p63γ-associated EMT is promoted through a *TP63*/*SRC*/*SNAIL2* transcriptional axis (Srivastava et al. 2018). However in HNSC patients, we failed to observe any increased accumulation of SRC in TCGA tumor samples with higher expression of the gamma isoform (data not shown). Another possibility arises from the existing feed-back loop where Wnt/B-catenin signaling promotes *TP63* expression (Haas et al. 2019; Ruptier et al. 2011) and p63 represses Wnt/B-catenin signaling (Katoh et al. 2016; Lakshmanachetty et al. 2018). This feedback loop could be essential for the maintenance of proper balance between differentiation and proliferation. One can speculate that this balance could be tilted by the modification of the activity of the p63 transcription factor associated with an altered ratio of the p63 C-terminal isoforms.

We observed that increased expression of *TP63γ* in tumors is associated with HNSC patients having a lower survival. While we can not conclude that this lower survival is a direct consequence of *TP63γ* expression, we identified PTBP1 as being more lowly in these tumor samples. It is therefore possible that the health status of these patients is associated with a PTBP1-specific altered splicing pattern that is reflected in the splicing of *TP63*. In this context measuring *TP63* isoforms in tumor biopsies could serve as a biomarker to identify patients with altered splicing patterns and potentially altered survival.

## Supporting information

reagents

## Acknowledgment

This work was funded by grants from Ligue Régionale contre le cancer (comité 22, 29, 35). WT was financed by a Contrat Doctoral from the Université de Rennes I. The authors wish to thank Christophe Tascon (IGDR-protein purification and analysis) for purification of the hsPTBP1 protein and Tiffany Yen Kway for her help with the correlation script.

