## supplementary_figures for "THE SPLICING FACTOR PTBP1 REPRESSES *TP63 γ* ISOFORM PRODUCTION IN SQUAMOUS CELL CARCINOMA"

### SUPPLEMENTARY FIGURE 1

**A** Correspondance between the TP63 gene model from TCGA and GTEX

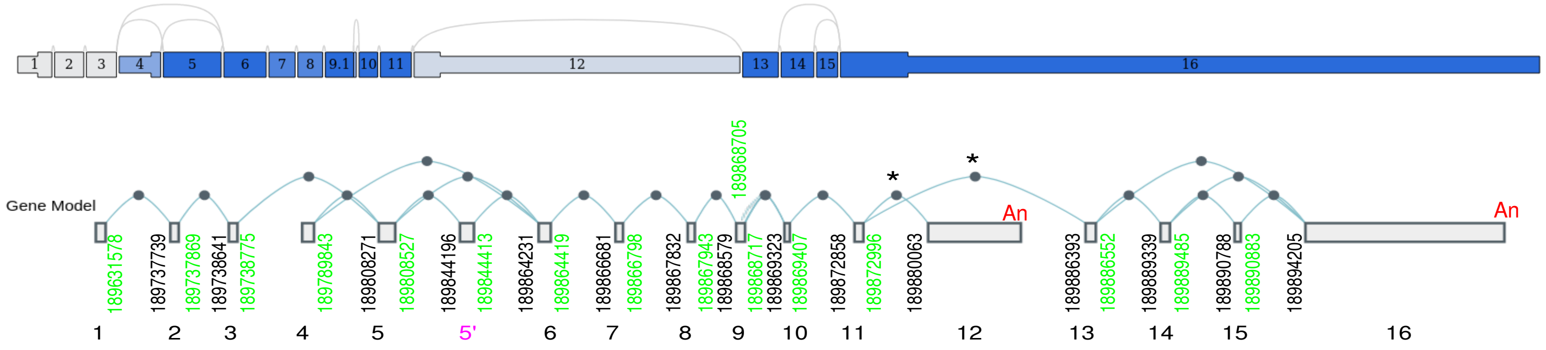


**B** Exon composition of the different C-terminal isoforms

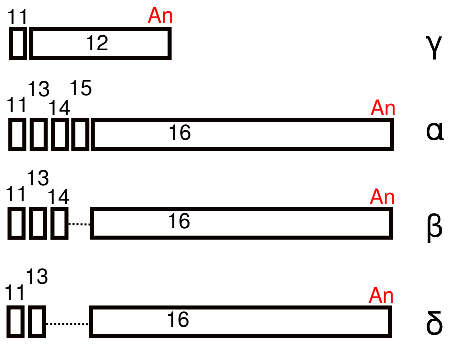

### SUPPLEMENTARY FIGURE 2

**A** TP63 gene structure - 21 junctions

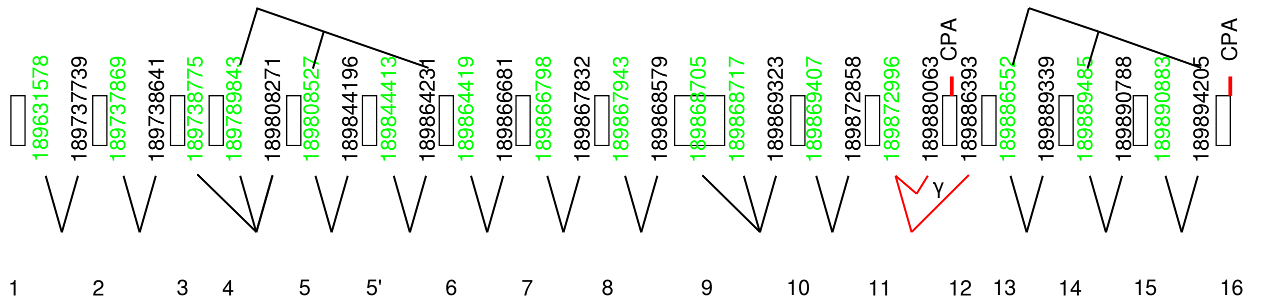


**B** TP63 Junction usage in TP63 expressing tissues

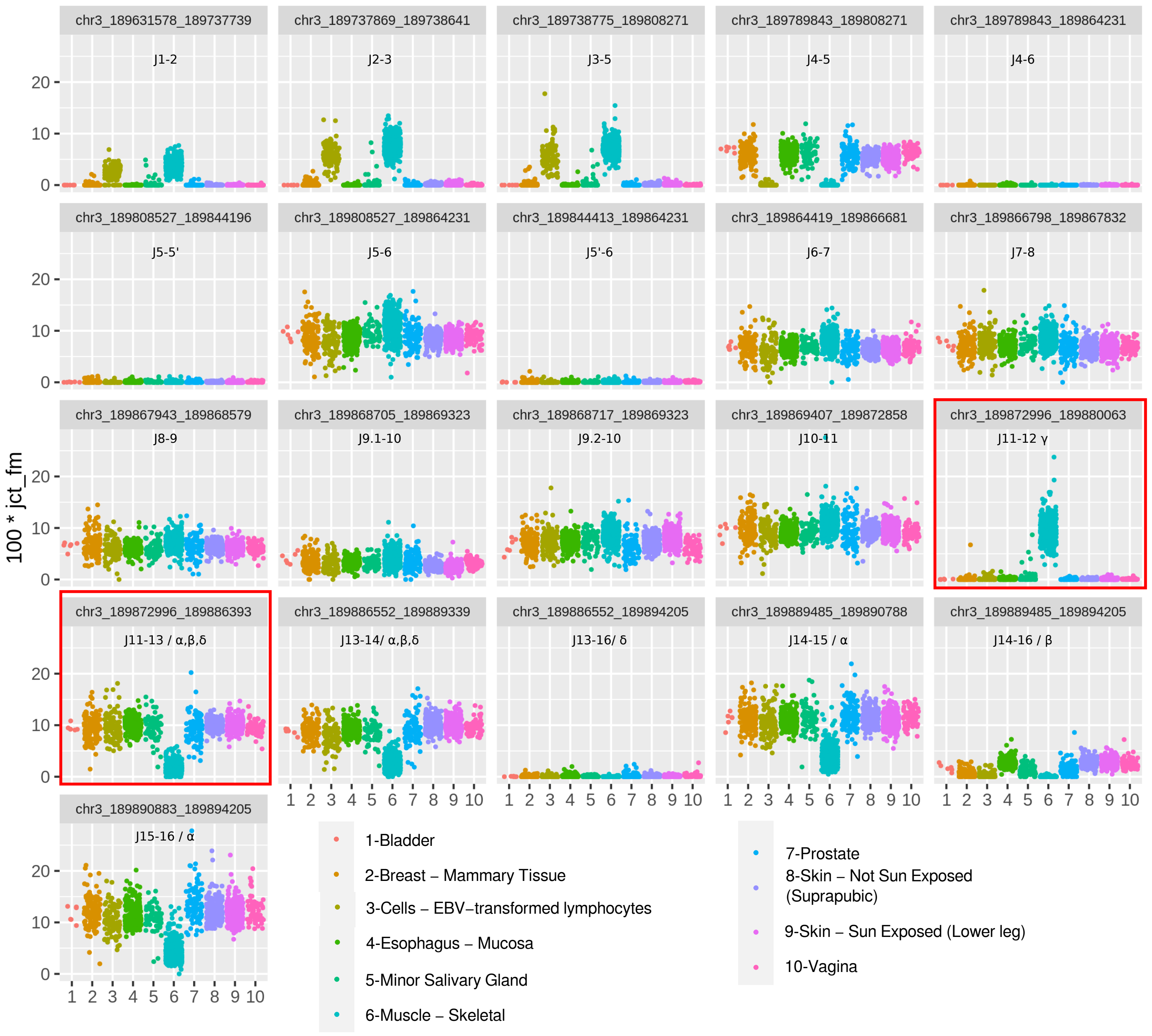

#### Supplementary Figure 3

**A**

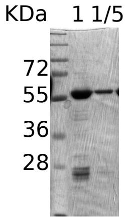


hs_Ptbp1_HIS6
is 60.5 KDa

**B**

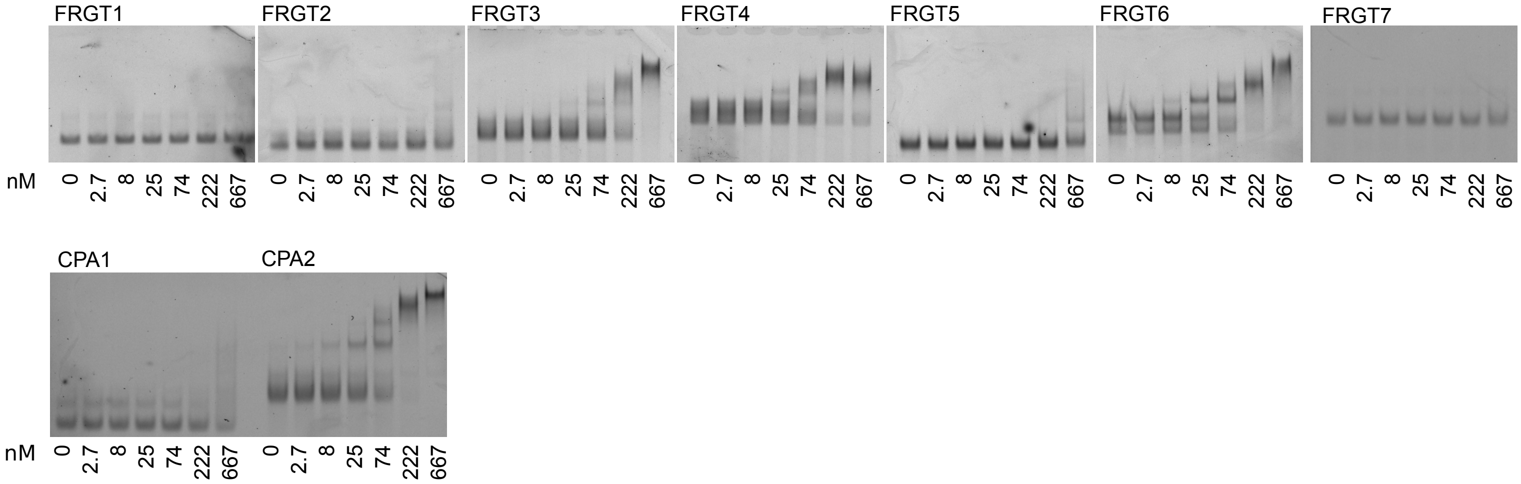

#### SUPPLEMENTARY FIGURES

##### Supplementary Figure 1: TP63 gene structure

A) Correspondence of the TCGA (upper lane) and GTEX (lower lane) annotations for the *TP63* locus. Hg38 chromosomal locations of 5’ and 3’ splice sites are shown in green and black respectively. The main difference between both annotations stands in the presence of an additional exon, noted 5’ here, in the GTEX gene model. The exon numbering used throughout the manuscript is from the TCGA annotation and presented underneath the GTEX gene model. B) Exon composition of the C-terminal isoforms γ, α, β, δ.

##### Supplementary Figure 2: Alternative junction usage for TP63 in GTEX data

A) Schematic of the TP63 gene model. The exons (boxes) and the quantified splice junctions (lines) are presented with their hg38 coordinates (green 5’ splice sites, black 3’ splice sites). Red ticks indicate cleavage and polyadenylation sites (CPA). The junctions pertaining to isoform γ are shown in red. B) junction usage among GTEX samples for the *TP63* expressing tissues. The genomic coordinates of the junctions, the exons joined and the isoform involved (when specific) are shown on each graph.

##### Supplementary Figure 3: Electrophoretic mobility shift assays

A) Validation of the purification of the recombinant human PTBP1 protein by gel electrophoresis and coomassie staing. B) Representative EMSA experiments conducted with fluorescently labeled RNA and recombinant PTBP1 proteins in increasing concentration (0 to 667 nM) (n >= 4). In each panel the lower bands in 0 nM PTBP1 correspond to free RNA probes.
